# Longitudinal tracking of *Plasmodium falciparum* clones in complex infections by amplicon deep sequencing

**DOI:** 10.1101/306860

**Authors:** Anita Lerch, Cristian Koepfli, Natalie E. Hofmann, Johanna H. Kattenberg, Anna Rosanas-Urgell, Inoni Betuela, Ivo Mueller, Ingrid Felger

## Abstract

**Background:** Longitudinal tracking of individual *Plasmodium falciparum* strains in multi-clonal infections is essential for investigating infection dynamics of malaria. The traditional genotyping techniques did not permit tracking changes in individual clone density during persistent natural infections. Amplicon deep sequencing (Amp-Seq) offers a tool to address this knowledge gap.

**Methods:** The sensitivity of Amp-Seq for relative quantification of clones was investigated using three molecular markers, *ama1-D2, ama1-D3,* and *cpmp.* Amp-Seq and length-polymorphism based genotyping were compared for their performance in following minority clones in longitudinal samples from Papua New Guinea.

**Results:** Amp-Seq markers were superior to length-polymorphic marker *msp2* in detecting minority clones (sensitivity Amp-Seq: 95%, *msp2:* 85%). Multiplicity of infection (MOI) by Amp-Seq was 2.32 versus 1.73 for *msp2.* The higher sensitivity had no effect on estimates of force of infection because missed minority clones were detected in preceding or succeeding bleeds. Individual clone densities were tracked longitudinally by Amp-Seq despite MOI>1, thus providing an additional parameter for investigating malaria infection dynamics.

**Conclusion:** Amp-Seq based genotyping of longitudinal samples improves detection of minority clones and estimates of MOI. Amp-Seq permits tracking of clone density over time to study clone competition or the dynamics of specific, i.e. resistance-associated genotypes.

## Introduction

Molecular-epidemiological parameters used to describe the infection dynamics of *Plasmodium falciparum* include the number of co-infecting parasite clones (multiplicity of infection, MOI), the rate at which different genotypes are acquired over time (molecular force of infection,_mol_FOI) and duration of infection^1^. These measures are based on monitoring the presence or absence of clones in cross-sectional or longitudinal samples collected in regular intervals. In earlier studies individual parasite clones in multi-clonal field samples were distinguished and tracked over time by genotyping the length-polymorphic marker merozoite surface protein 2 *(msp2)* by capillary electrophoresis-based fragment sizing (CE)^2–4^. Yet, *msp2*-CE genotyping has limited sensitivity for minority clone detection^3,5^. Alternative typing methods instead could perform better in detecting minority clones, but might impact measures of MOI and _mol_FOI^6^. So far quantification of individual clones within multi-clonal infections was not feasible, as this would have required highly complex allele-specific quantitative PCR (qPCR).

SNP-based genotyping by deep amplicon sequencing (Amp-Seq) can detect low-abundant *P. falciparum* clones at ratios of 1:1000 in mixed infections^7^. Most importantly, genotyping by Amp-Seq also quantifies precisely the relative abundance of clones, as shown with artificial mixtures of clones^7–9^. From these ratios the absolute density of each clone (i.e. a certain haplotype) within a multi-clone infection can be deduced if the total parasitaemia of the sample was established by qPCR^9^. When analysing consecutive samples from a given study participant, presence and fluctuations in density of clones can be tracked. We explore how longitudinal information can be used to improve identification of minority clones with low densities around the detection limit.

A previous study has estimated clonal density with Amp-Seq in multi-clone infections to estimate clearance rates after antimalarial treatment^9^. We apply the same approach to track parasite clones longitudinally in untreated natural infections. In addition, we increase the resolution of genotyping by combining sequence information from several markers into multi-locus haplotypes.

## Methods

### Study design

A subset of 153 archived *P. falciparum* genomic DNA samples from 33 children (mean 4.3 samples [min: 2, max: 11]) aged 1-5 years were available from an cohort study with blood sampling over 40 weeks (first 12 weeks every fortnightly, then monthly) in Papua New Guinea (PNG)^10^. The two conditions for selection of children were: ≥2/14 bleeds PCR positive, and MOI>1 in at least one of the samples of each child. Ethical clearance was obtained from PNG Institute of Medical Research Institutional Review Board (IRB 07.20) and PNG Medical Advisory Committee (07.34). Informed written consent was obtained from all parents or guardians prior to recruitment of each child.

### Genotyping using length polymorphic marker *msp2*

Samples were genotyped using the classical *P. falciparum* marker *msp2* according to published protocols^11^. Fluorescently labelled nested PCR products were sized by CE on an automated sequencer and analysed using GeneMarker software. Fragments were accepted if the following cut-off criteria were met: peak height >500 intensity units and >10% of the height of the majority peak. Electropherograms were inspected visually to exclude obvious stutter peaks. All DNA samples were genotyped in 2 independent laboratories to assess reproducibility of clone detection and measures of MOI.

### Marker selection for Amplicon deep sequencing

Amp-Seq was performed on three amplicons located in two different *P. falciparum* marker genes, namely PF3D7_0104100, “conserved *Plasmodium* membrane protein” (cpmp), and PF3D7_1133400, “apical membrane antigen 1” *(ama1)* whose genetic diversity has been studied in great detail^12–14^. Previously published primers were used for marker *cpmp*^7^. For *ama1* two amplicons of 479 and 516 bp were selected that span regions of maximum diversity, i. e. subdomains 2 and 3 of the ectodomain^15^. Primer sequences and exact amplicon positions are listed in Supplementary Tables S1 and S2.

### Sequencing library preparation

Sequencing libraries were generated by three rounds of PCR, according to previously published protocols^7^. After primary PCR, a 5’ linker sequence was added during nested PCR. Nested PCR products were subject to another PCR round with primers binding to the linker sequences and carrying Illumina sequence adapters plus an eight nucleotide long sample-specific molecular index to permit pooling of amplicons for sequencing and later de-multiplexing. The final sequence library was purified with NucleoMag beads prior to sequencing on an Illumina MiSeq platform in paired-end mode using Illumina MiSeq reagent kit v2 (500-cycles) together with Enterobacteria phage PhiX control (Illumina, PhiXControl v3).

### Sequence read analysis and haplotype calling

Samples yielding a sequence coverage of <25 reads were excluded from the analysis. An overview of sequence read coverage for all Amp-Seq markers is given in Supplementary Table S3. Sequence reads were analysed using software HaplotypR (https://github.com/lerch-a/HaplotypR.git). Haplotype calling is explained in full detail in an earlier publication^7^. In short: Low quality sequences were removed by trimming forward reads to 240bp and reverse reads to 170bp. As reference sequence *P. falciparum* strain 3D7 was used (PlasmoDB release 34,^16^). The term genotype refers to a single nucleotide polymorphism (SNP). Calling a SNP required a >50% mismatch rate in the sequence reads of this nucleotide position in at least two independent samples. A haplotype was defined as sequence variant of an entire amplicon. Haplotypes containing inserts or deletions (indels) were filtered out, as well as haplotypes resulting from chimeric reads or singleton reads. The number of reads of a given haplotype over all remaining reads of the same marker within a sample is denoted by the term “within-host haplotype frequency”. Cut-off criteria for haplotype calling were as follows: a minimum of 3 reads coverage per sample, a within-host haplotype frequency ≥0.1% and an occurrence of this haplotype in at least 2 samples.

### Multi-locus haplotype inference in longitudinal samples

Amp-Seq quantifies the frequency of each haplotype within a sample, which permits to infer multi-locus haplotypes. A multi-locus haplotype was deduced in multiple rounds. In the first round, the multi-locus haplotype of the dominant clone of a sample was inferred by selecting each marker’s dominant haplotype (>54% within-host haplotype frequency, i.e. 50%+3.8% standard deviation in within-host haplotype frequency between replicates). After each round the identified dominant haplotype was ignored and in the following round the dominant haplotype was identified among the remaining reads. If several haplotypes occurred in a sample at similar frequencies, it may be impossible to identify the dominant haplotype. This was resolved by analysing the change in within-host haplotype frequency between the observed and preceding or succeeding sample of the same host. An example of our approach to multi-locus haplotype inference is shown in detail the Supplementary Information S1.

The final step of multi-locus haplotype inference addressed the problem of clones of a multiple infection that share by chance the same allele of one of the markers. As a consequence, the within-host frequency of a shared haplotype amounts to the sum of two or more independent clones carrying the same allele. In such cases multi-locus haplotypes were inferred by assigning the shared alleles to those haplotypes that summed up to the same proportion in the other two markers. Samples for which the multi-locus haplotype could not be established by this approach were considered unresolvable (Supplementary Table S4).

### Reproducibility, sensitivity and false discovery rate

Samples were analysed in duplicates with Amp-Seq markers and *msp2*-CE. Performing duplicates permitted to identify and exclude false-positive haplotypes and thus prevented erroneous over-estimation of MOI. Each haplotype was classified into one of four groups (example see Supplementary Figure S1): (1) True-positive (TP) haplotype, i.e. it passed the haplotype calling cut-off in both replicates or in one replicate plus in the preceding or succeeding bleed; (2) False-positive (FP) haplotype, i.e. it passed the haplotype calling cut-off in only one replicate and was not detected in any of the preceding or succeeding samples of that individual; (3) False-negative (FN_i_) haplotype, i.e. it was detected in one or both replicates but did not pass the cut-off criteria at that occasion, whereas it was detected in the preceding or succeeding bleed as TP (at least once) or FN haplotype; (4) Background noise (all other cases).

Additionally, false-negative (FN_ii_) haplotypes were imputed for samples in which no sequence read was detected. These false-negative haplotypes were imputed only when (a) the haplotype was detected in the preceding as well as the succeeding bleed as a true-positive. Presence in only one of preceding or succeeding sample was not considered sufficient evidence for assuming a case of missed detection. For the Amp-Seq markers but not *msp2*-CE, false-negative haplotypes were also imputed when (b) data for the other two markers was present and the corresponding multi-locus haplotype was established in the preceding or succeeding sample.

The sensitivity to detect parasite clones was estimated based on selected individuals who had not received antimalarial treatment during the timespan analysed and harboured at least one haplotype that was detected at 3 consecutive bleeds. Sensitivity was defined as the true positive rate of a genotyping method and was calculated as TP/(TP+FN). The risk to falsely assign a haplotype not present in the sample was measured as the “false discovery rate” (FDR), calculated as FP/(TP+FP). This rate represents the extent of false haplotype calls of a genotyping method.

The reproducibility of clone detection in technical replicates (comprising all experiential procedures from PCR to sequence run) was calculated as 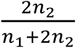, where *n*_1_ is the number of haplotypes detected in a single replicate and *n*_2_ the number of haplotypes detected in both replicates^17^. Only TP haplotypes were used to estimate reproducibility.

### Epidemiological parameters: clone density, diversity, MOI and FOI

The density of a parasite clone was calculated by multiplying within-host haplotype frequency by parasitaemia (measured by qPCR). Clone density is expressed as copies of target gene per microliter, quantified by qPCR targeting the 18S rRNA gene of *P. falciparum*^18^. The technical detection limit of qPCR was 0.4 copies/μl whole blood.

Based on true positive haplotypes, the expected heterozygosity (H_e_) and mean MOI were determined from baseline (or first bleed available) samples for each marker as described^7^. H_e_ was also estimated for combined markers in samples that had a resolvable multi-locus haplotype and that were separated by a treatment plus ≥2 consecutive *P. falciparum* negative samples from the same child.

_mo1_FOI was estimated on longitudinal sets of sample that had a complete set of replicates. Haplotypes were counted as new infection if a haplotype was (i) not present in the baseline sample but in a subsequent sample, (ii) not detected at ≥2 consecutive preceding bleeds or (iii) not detected after antimalarial treatment plus after at least one negative sample. Time at risk was calculated as the timespan between baseline and last sampling, minus 14 days for each antimalarial treatment (to account for the prophylactic effect of treatment).

An overview of sample selection criteria applied for different types of analyses is listed in Supplementary Table S5.

## Results

### Genetic diversity of markers

The discriminatory power of Amp-Seq markers *cpmp, ama1-D2* and ama1-D3, as well as length-polymorphic marker *msp2*-CE was estimated in 33 baseline samples. The resolution was highest for amplicon marker *cpmp* (H_e_=0.961) that distinguished 30 haplotypes and gave a mean MOI=2.45 (Table 1, MOI distribution by marker in Supplementary Figure S2). The second-best resolution was obtained by marker *msp2*-CE (H_e_=0.940) that distinguished 20 haplotypes and measured a mean MOI=1.73. Haplotype and SNP frequencies of Amp-Seq markers are shown in Figure 1 and Supplementary Figure S2.

**Figure 1:**
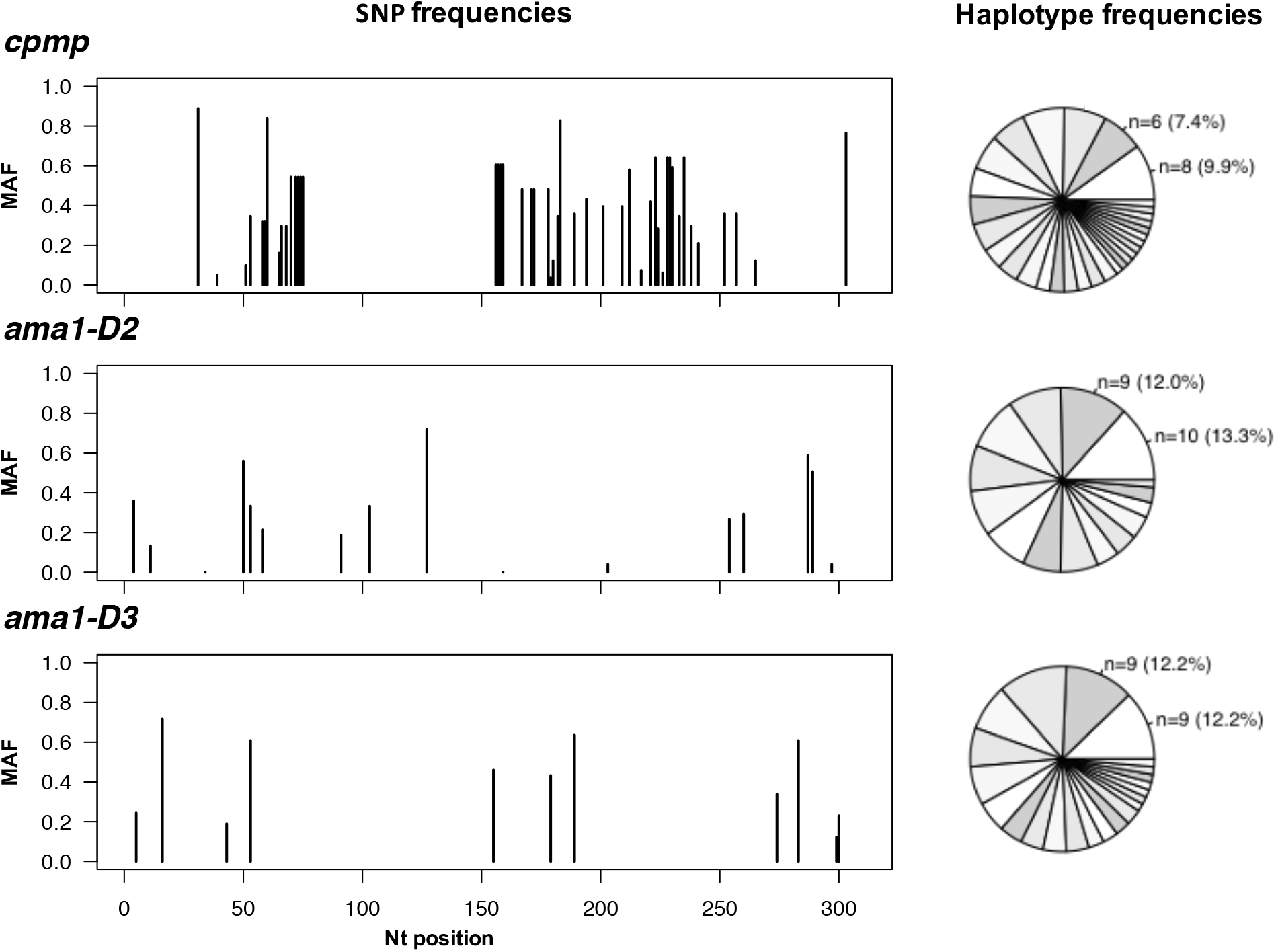
Frequency of individual SNPs and haplotypes of three markers in 33 baseline samples from PNG. Minor allelic frequency (MAF) of each SNP (left) and frequency of haplotypes in these baseline samples (right), n, number of observations per haplotype shown for 2 most prevalent haplotypes. Total number of different haplotypes: 30 for *cpmp,* 15 for *ama4-*D2 and 22 for *ama4-*D3. (Frequency of haplotypes for markers *msp2*-CE given in Supplementary Figure S2).

**Table 1:** Genotyping results of 4 molecular markers analysed in 33 baseline field samples. H_e_, expected heterozygosity. MOI, multiplicity of infection.

| Marker | $H_e$ | Mean MOI | Number of clones<sup>1</sup> | Number of haplotypes | Number of SNPs<sup>2</sup> |
| --- | --- | --- | --- | --- | --- |
| *msp2* CE | 0.940 | 1.73<sup>3</sup> | 57 | 20 | n/a |
| *cpmp* | 0.961 | 2.45<sup>3</sup> | 81 | 30 | 48 |
| *ama1*-D2 | 0.928 | 2.27<sup>3</sup> | 75 | 15 | 17 |
| *ama1*-D3 | 0.939 | 2.24<sup>3</sup> | 74 | 22 | 11 |

<sup>1</sup> Sum of all haplotypes in all samples.
<sup>2</sup> With respect to the reference sequence of *P. falciparum* strain 3D7.
<sup>3</sup> Pairwise comparison using two-sided paired t-test with adjusted p-value by Holm: p-value=0.008 for *ama1*-D2 vs *msp2*-CE, p-value=0.036 for *ama1*-D3 vs *msp2*-CE, and p-value=0.005 for *cpmp* vs *msp2*-CE.

Discriminatory power can be increased by combining multiple markers. Inference of multi-locus haplotypes was possible for 66 clones in 46 selected samples. Combining marker *cpmp* with either of the two *ama1* fragments yielded very high diversity (53 and 55 haplotypes, H_e_=0.992 and 0.994 for *cpmp/ama1-D2* and cpmp/ama1-D3) (Table 2 and Supplementary Figure S3). Combining all 3 markers did not increase discriminatory power any further.

**Table 2:** Genotyping results of 3 molecular markers analysed in 47 independent field samples with 66 different clones. H_e_, expected heterozygosity.

| **Marker** | **$H_e$** | **Number of**<br>**Haplotypes** |
| --- | --- | --- |
| *cpmp* | 0.948 | 25 |
| *ama1-D2* | 0.926 | 16 |
| *ama1-D3* | 0.938 | 21 |
| *cpmp* + *ama1-D2* | 0.992 | 53 |
| *cpmp* + *ama1-D3* | 0.994 | 55 |
| *cpmp* + *ama1-D2* + *ama1-D3* | 0.994 | 55 |

### Using longitudinal genotyping data to increase detectability of clones

Imperfect detectability of parasite clones has been described previously in longitudinal genotyping studies^1,19–21^. Data from replicates and longitudinal samples can be used to make assumptions on missed clones. This permits imputing of missed haplotypes and thus improves the tracking of clonal infections within an individual over time. Two types of missed haplotypes respective false-negative haplotypes were distinguished: (FN_i_) haplotypes that were detected below the cut-off and (FN_ii_) haplotypes that were not detected but imputed (Supplementary Table S6). Supplementary Figure S4 shows an example of different type of missed haplotypes for all Amp-Seq markers.

The sensitivity to detect parasite clones was estimated for each genotyping marker by enumerating false-negative haplotypes. Sensitivity was higher for the Amp-Seq markers than for *msp2*-CE (in decreasing order 96.5%, 95.0%, 93.9% and 85.1% for *ama1-D2, cpmp, ama1-D3* and *msp2*-CE) (Table 3). For ≥57% of the identified false-negative haplotypes, reads were detected but fell below cut-off criteria (category (i) above). If such haplotypes were counted as positives by relaxing the cut-off criteria, sensitivity would increase to 99.1%, 97.5% and 97.4% for Amp-Seq markers ama1-D2, *cpmp* and ama1-D3 (Table 3).

**Table 3:**
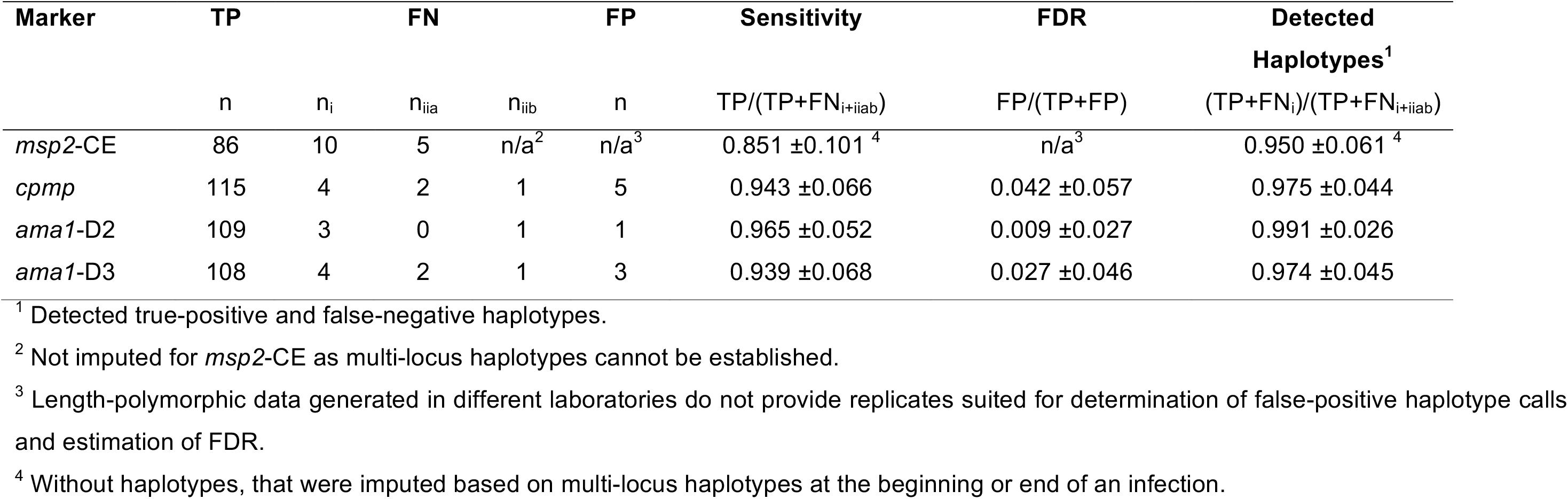
Sensitivity and false discovery rate (FDR) of the genotyping method. Sensitivity and FDR including 95% confidence interval was estimated based on persistent clones in 48 longitudinal samples from 12 individuals. Detectability of minority clone can be increased by including missed persistent haplotypes detected below the cut-off criteria. TP, true-positive haplotypes. FN_i_, false-negative haplotypes detected, but below cut-off criteria. FN_iiab_, false-negative haplotypes with no read detected.

The false discovery rate of haplotypes for Amp-Seq markers was in the range of 0.9-4.2% (Table 3). Reproducibility to detect parasite clones in technical replicates was greater for Amp-Seq markers than for marker *msp2*-CE (in decreasing order 0.95, 0.95, 0.94 and 0.91 for *ama1-D3, cpmp, ama1-D2* and *msp2*-CE) (Supplementary Table S7 and Figure S5).

### Determination of _mol_FOI by different molecular markers and methods

A higher sensitivity of the genotyping method does not necessary impact molFOI, i.e. new clones/year, because a missed minority clone could be detected at one of the successive bleeds. We investigated the number of new infections acquired during 40 weeks follow-up in 27 children from whom a complete data set was available (on average 4.3 samples per child [min: 2, max: 7]). Mean molFOI was 2.7, 2.7, 2.3 and 2.2 new infections per year for markers *ama4-* D3, *cpmp, msp2*-CE and *ama1*-D2 (negative binomial regression p-value for comparison of *msp2*-CE to *ama1*-D3, *cpmp* and *ama1*-D2: 0.596, 0.649 and 0.877) (Supplementary Figure S6). Thus, no substantial difference in mean _mol_FOI was found for the different molecular markers and different genotyping methods.

### Quantitative dynamics of multiple infecting *P. falciparum* clones

Densities of individual clones was calculated from the total parasitaemia by qPCR and the within-host haplotype frequency. Examples of individual clone density dynamics in children with multi-clone infections are shown for three Amp-Seq markers (Figure 2). The density of some clones remained constant over time, whereas other clones showed fluctuations in density over 3 orders of magnitude (Figure 2A and B). In some children the dominant clone remains dominant over the observation period (Figure 2A), whereas in others switch-over between minority clone and dominant clone was observed (Figure 2B). In highly complex field samples some clones might share the same haplotype of a given marker (Figure 2C). Such clones can only be differentiated and quantified if multiple markers are typed and at least one of the markers is not shared between concurrent clones.

**Figure 2:**
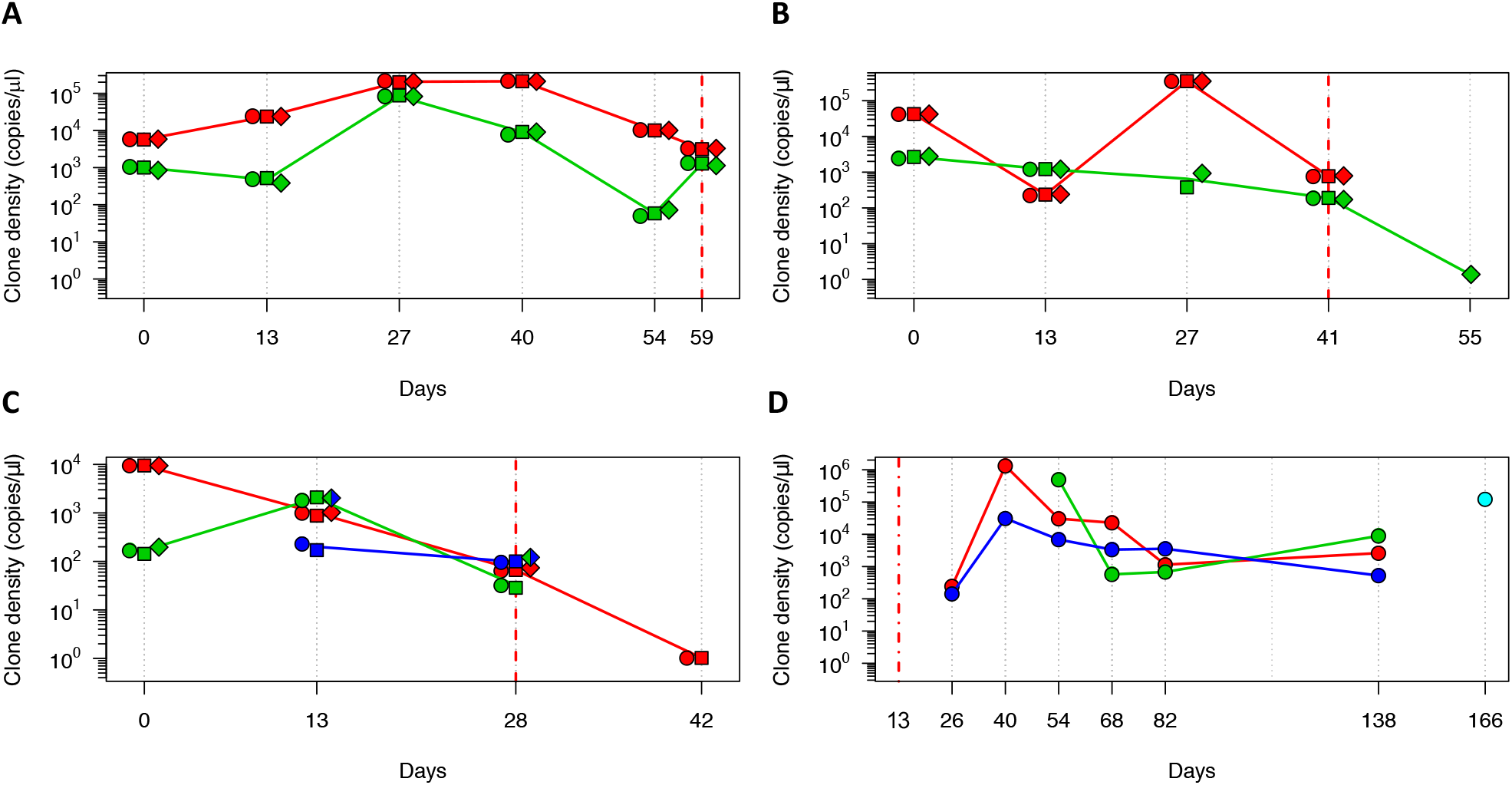
Dynamics of multi-clone infections in 4 children. Multi-marker haplotypes could be generated in panels A, B and C. Inference of multi-locus haplotypes was not possible for the child in panel D; here the dynamics of individual clones tracked by marker ama1-D2 are shown. Each colour represents a clone, individual markers represented by different shapes: *cpmp* (diamonds), *ama1-*D2 (circles) and *ama1-*D3 (squares). Solid line connecting multi-locus haplotypes represents their median frequency. Grey dotted vertical lines represent sampling dates. Red dashed lines represent day of artemisinin combination therapy. Red dash-dotted line represents end of radical cure (artesunate-primaquine) at baseline.

After artemisinin combination therapy, some of the parasite clones from multi-clone infections were cleared 14 days after antimalarial treatment, whereas others were still detectable (Figure 2A, B and C). These persisting clones had decreased clone densities (<21 copies/μl) and likely represent remaining late gametocyte stages of cleared asexual infections^22^. Some new infections following antimalarial treatment (artesunate-primaquine) showed a rapid increase in clone density within the first 14 days after re-infection of a host, followed by a slow decrease in clone density until clearance (Figure 2D), whereas in other infections clone density remained constant (Figure 2C).

## Discussion

While MOI and _mol_FOI have been extensively described as epidemiological parameters, the ratio and density of individual clones within complex infections has not yet been investigated. This gap in knowledge was due to shortfalls of traditional length-polymorphic markers, where the length of a fragment greatly influences the amplification efficiency in multi-clone infections with fragments competing in PCR and a strong bias favouring smaller fragments^5^. As a result, multilocus haplotypes could not be inferred from traditional genotyping data in a reliable way. Such inference is required, for example, for phylogenetic or population genetic studies. In such studies, multiple-clone infections were usually excluded or only the predominant haplotype included^23,24^. With the possibility to establish multi-locus haplotypes from complex infections the discriminatory power will be greatly improved in future. This study explored the feasibility of multi-locus haplotype calling in complex infections and the usefulness of the Amp-Seq genotyping technique in longitudinal data.

Single Amp-Seq markers *cpmp*, ama1-D2, ama1-D3, and *msp2*-CE yielded similar resolution. Combining *cpmp* with either of the *ama1* fragments increased further discriminatory power. The excellent performance of Amp-Seq marker *cpmp* had been demonstrated earlier^7^. Such increased resolution is of great practical value for PCR-correction in clinical drug efficacy trials, where new infections need to be reliably distinguished from those present in an individual earlier. Robust methods for this application are urgently needed.

For infections with high multiplicity (MOI≥3), inference of multi-locus haplotypes remains challenging (example in Supplementary Table S7). Inference is straightforward if haplotypes occur at distinctive abundance in any of the longitudinal samples. If haplotypes are equally abundant in a sample and remain so over time, the multi-locus haplotype cannot be inferred. The same is true for complex patterns of shared haplotypes. In the present study, multi-locus haplotypes up to MOI=3 were inferred. For higher multiplicity, sophisticated statistical methods like Markov chain Monte Carlo on longitudinal samples could be applied^25^.

Genotyping longitudinal samples in duplicates enabled an evidence-based approach to identify false-negative haplotypes. This permitted to estimate each marker’s sensitivity to detect minority clones. Amp-Seq genotyping with markers *ama1-D2, ama1-D3* and *cpmp* missed less clones compared to *msp2*-CE genotyping (Amp-Seq in average 5.4% versus 14.9% *msp2*-CE). This difference is likely due to less stringent cut-off criteria for Amp-Seq compared to *msp2* genotyping. Minority clone detection by *msp2*-CE is limited by peak calling cut-off criteria, which are usually a fixed minimal signal intensity plus a minimum peak height of 10% (used in our study) or more of the dominant peak. Minority clones with an abundance of <10% of all amplified fragments will not pass these criteria. An increase of *msp2*-CE sensitivity would require a lower cut-off, which would lead to more false positive signals from either stutter peaks or background noise. In contrast, Amp-Seq allows to remove PCR artefacts before haplotype calling and thus can support a much lower cut-off of <1%^7^.

In cohort studies where Amp-Seq genotyping is performed in successive follow up samples of the same patient, an even more relaxed definition of Amp-Seq cut-off criteria would be justifiable. In this scenario, the same evidence-based strategy of using successive samples can be used to recover minority haplotypes that were detected with read counts below the haplotype calling cut-offs. If recovery would be performed in this study, ≥57% of all false-negative haplotypes would be identified. Such recovery would increase detectability of parasite clones by Amp-Seq to >97%. In addition, multi-locus haplotypes could provide additional evidence for accurate recovery.

The higher sensitivity of Amp-Seq to detect minority clones compared to *msp2-CE* substantially increased MOI, but did not affect mean _mol_FOI. Any estimation of _mol_FOI needs to account for temporary absence of clones from the peripheral blood caused by sequestration^1,19–21^. A clone that is temporarily undetectable owing to density fluctuations is likely observed at either the preceding or succeeding bleed. Therefore, a clone is usually only counted as new infection if it was not detected in ≥2 consecutive blood samples. As a consequence, a clone missed at a single bleed will not necessarily lead to a decrease of molFOI.

A clone that was intermittently missed at one bleed by *msp2-CE* was always detected by Amp-Seq. This observation supports the practice in earlier papers where intermittently missed clones were imputed^21^. Counting a recurrent haplotype as new infection after a single negative bleed would lead to an overestimation of _mol_FOI^1,19–21^. The statistical power of this study was limited and a larger study is needed to fully explore the effect of the typing method used on estimates of MOI, molFOI, or even prevalence rates.

A major advantage of Amp-Seq over *msp2*-CE is that the density of an individual clone in multiclone infections can be calculated. Quantifying the density of individual parasites clones over time permits to study dynamics, and thus fitness, of parasite clones exposed to within-host competition^26^. For example, the relative densities of new infections can be compared to clones already persisting in a host, and their densities in respect to extrinsic factors or clinical symptoms can be investigated.

## Conclusion

Amplicon sequencing improves clone detectability compared to *msp2*-CE owing to its greater sensitivity for detection of minority clones. Our results confirm earlier assumptions on clone persistence with intermittent missed observation. This validates the imputation of false negatives to correct for imperfect detection of clones, a strategy also used in previous studies on clone dynamics. Using multi-locus haplotypes for genotyping permitted to identify robustly individual clones and improved differentiation between new and recurring clones. Construction of multilocus haplotypes are of great value to compensate the effects of highly abundant haplotypes in the population. The option to quantify individual clones enables new approaches to investigate effects of parasite fitness or superinfection in multi-clone infections.

## Acknowledgement

We are grateful to the study participants and their guardians and to the field and laboratory team of the PNG Institute of Medical Research, in particular Alice Ura. We would also like to thank Stephen Wilcox for supervising sequence library preparation and sequencing.

## Authors Contributions

Conceived and designed the experiments: IF, IM, AL, CK. Performed the experiments: AL, CK, JHK, NH, ARU. Supervised field work: IB, ARU. Analysed the data: AL. Supervision: IF. Writing-draft: AL, IF. All Co-authors have read the manuscript and agreed with the final version.

## Reference

1. Felger, I. et al. The Dynamics of Natural Plasmodium falciparum Infections. PLoS One 7, e45542 (2012).

2. Hofmann, N. E. et al. The complex relationship of exposure to new Plasmodium infections and incidence of clinical malaria in Papua New Guinea. Elife 6, 1–23 (2017).

3. Koepfli, C. et al. How much remains undetected? Probability of molecular detection of human Plasmodia in the field. PLoS One 6, e19010 (2011).

4. Sonden, K. et al. Asymptomatic Multiclonal Plasmodium falciparum Infections Carried Through the Dry Season Predict Protection Against Subsequent Clinical Malaria. J. Infect. Dis. 212, 608–616 (2015).

5. Messerli, C., Hofmann, N. E., Beck, H.-P. & Felger, I. Critical Evaluation of Molecular Monitoring in Malaria Drug Efficacy Trials and Pitfalls of Length-Polymorphic Markers. Antimicrob. Agents Chemother. 61, AAC.01500-16 (2017).

6. Miller, R. H. et al. A deep sequencing approach to estimate Plasmodium falciparum complexity of infection (COI) and explore apical membrane antigen 1 diversity. Malar. J. 16, 490 (2017).

7. Lerch, A. et al. Development of amplicon deep sequencing markers and data analysis pipeline for genotyping multi-clonal malaria infections. BMC Genomics 18, 864 (2017).

8. Levitt, B. et al. Overlap Extension Barcoding for the Next Generation Sequencing and Genotyping of Plasmodium falciparum in Individual Patients in Western Kenya. Sci. Rep. 7, 41108 (2017).

9. Mideo, N. et al. A deep sequencing tool for partitioning clearance rates following antimalarial treatment in polyclonal infections. Evol. Med. public Heal. 2016, 21–36 (2016).

10. Betuela, I. et al. Relapses contribute significantly to the risk of Plasmodium vivax infection and disease in Papua New Guinean children 1-5 years of age. J. Infect. Dis. 206, 1771–80 (2012).

11. Falk, N. et al. Comparison of PCR-RFLP and Genescan-based genotyping for analyzing infection dynamics of Plasmodium falciparum. Am. J. Trop. Med. Hyg. 74, 944–50 (2006).

12. Arnott, A. et al. Distinct patterns of diversity, population structure and evolution in the AMA1 genes of sympatric Plasmodium falciparum and Plasmodium vivax populations of Papua New Guinea from an area of similarly high transmission. Malar. J. 13, 233 (2014).

13. Cortés, A. et al. Allele specificity of naturally acquired antibody responses against Plasmodium falciparum apical membrane antigen 1. Infect. Immun. 73, 422–30 (2005).

14. Cortés, A. et al. Geographical structure of diversity and differences between symptomatic and asymptomatic infections for Plasmodium falciparum vaccine candidate AMA1. Infect. Immun. 71, 1416–26 (2003).

15. Hodder, A. N. et al. The disulfide bond structure of *Plasmodium* apical membrane antigen-1. J. Biol. Chem. 271, 29446–52 (1996).

16. Bahl, A. et al. PlasmoDB: The Plasmodium genome resource. A database integrating experimental and computational data. Nucleic Acids Research 31, 212–215 (2003).

17. Bretscher, M. T. et al. Detectability of Plasmodium falciparum clones. Malar. J. 9, 234 (2010).

18. Rosanas-Urgell, A. et al. Comparison of diagnostic methods for the detection and quantification of the four sympatric Plasmodium species in field samples from Papua New Guinea. Malar. J. 9, 361 (2010).

19. Sama, W., Owusu-Agyei, S., Felger, I., Dietz, K. & Smith, T. Age and seasonal variation in the transition rates and detectability of Plasmodium falciparum malaria. Parasitology 132, 13–21 (2006).

20. Sama, W., Owusu-Agyei, S., Felger, I., Vounatsou, P. & Smith, T. An immigration-death model to estimate the duration of malaria infection when detectability of the parasite is imperfect. Stat. Med. 24, 3269–88 (2005).

21. Smith, T., Felger, I., Fraser-Hurt, N. & Beck, H. P. Effect of insecticide-treated bed nets on the dynamics of multiple Plasmodium falciparum infections. Trans. R. Soc. Trop. Med. Hyg. 93 Suppl 1, 53–7 (1999).

22. Bousema, T. et al. Revisiting the circulation time of Plasmodium falciparum gametocytes: molecular detection methods to estimate the duration of gametocyte carriage and the effect of gametocytocidal drugs. Malar. J. 9, 136 (2010).

23. MalariaGEN Plasmodium falciparum Community Project. Genomic epidemiology of artemisinin resistant malaria. Elife 5, 1–29 (2016).

24. Barry, A. E., Schultz, L., Buckee, C. O. & Reeder, J. C. Contrasting population structures of the genes encoding ten leading vaccine-candidate antigens of the human malaria parasite, Plasmodium falciparum. PLoS One 4, e8497 (2009).

25. Zhu, S. J., Almagro-Garcia, J. & McVean, G. Deconvolution of multiple infections in Plasmodium falciparum from high throughput sequencing data. Bioinformatics (2017). doi:10.1093/bioinformatics/btx530

26. de Roode, J. C., Culleton, R., Cheesman, S. J., Carter, R. & Read, A. F. Host heterogeneity is a determinant of competitive exclusion or coexistence in genetically diverse malaria infections. Proc. R. Soc. B Biol. Sci. 271, 1073–1080 (2004).

